# SLC25A51 impacts drug sensitivity in AML cells by sustaining mitochondrial oxidative flux

**DOI:** 10.1101/2022.11.15.516643

**Authors:** Mu-Jie Lu, Jonathan Busquets, Valeria Impedovo, Yu-Tai Chang, William Matsui, Stefano Tiziani, Xiaolu A. Cambronne

## Abstract

SLC25A51 imports oxidized NAD^+^ into the mitochondrial matrix and is required for sustaining oxidative metabolism in human mitochondria. We observed that higher expression of SLC25A51 correlated with poorer survival in Acute Myeloid Leukemia (AML) patient data. Given AML’s dependency on oxidative cell metabolism, we sought to determine the role SLC25A51 may serve in this disease. We found that depleting SLC25A51 in AML cells led to increased apoptosis, as well as prolonged survival in a xenograft model. Metabolic flux analyses indicated that depletion of SLC25A51 shunted flux away from oxidative pathways and promoted glutamine utilization for reductive carboxylation to support aspartate production. Consequently, SLC25A51 loss sensitized AML cells to glutamine deprivation and glutaminase inhibitor CB-839. Together, the work highlights connections between SLC25A51 and oxidative mitochondrial flux in AML. We identified a rationale for targeting SLC25A51 in myeloid cancers with potential for a therapeutic window, especially when coupled with glutaminase inhibition.

**Statement of significance:** This investigation describes an approach to directly modulate the tricarboxylic acid cycle as a potential vulnerability in oxidative tumors. Using AML models, the work is an inaugural look into SLC25A51’s role supporting oxidative mitochondrial metabolism and identifies SLC25A51 levels as a potential marker for stratification of AML.

## Introduction

Acute Myeloid Leukemia (AML) is an exemplar amid the increasing number of cancer types that depend on oxidative metabolism and a functional mitochondrial TCA cycle (1–4). Additionally, there is an unmet need to better understand this specific disease and for novel therapeutic approaches. AML occurs most frequently in older individuals for whom treatment options are limited, resulting in >70% mortality within the first five years due to refractory cases and relapsed disease (5).

The signature metabolism of AML has emerged as a promising vulnerability, and it includes a heightened reliance on oxidative metabolism, glutamine utilization, and oncometabolites (1–3). The challenge is that the contributions of each feature can vary across individual cases and change over time, thereby challenging our understanding of underlying mechanisms in specific cases at any given time. One shared characteristic, nevertheless, is that these identified vulnerabilities are all expected to depend on oxidized nicotinamide adenine dinucleotide (NAD^+^) in mitochondria. A direct study into the specific role of mitochondrial NAD^+^ in any cancer, however, has not been feasible due to the difficulty of targeting this compartment without affecting the whole cell.

Intracellular NAD^+^ is highly compartmentalized (6,7). It serves dual roles throughout the cell and in mitochondria. Through the transfer of hydride ions, NAD^+^ facilitates oxidative reactions, including those in the tricarboxylic acid (TCA) cycle. The relative levels of NAD^+^ compared to reduced NADH influence the local capacity for either oxidative or reductive reactions. Independently, mitochondrial NAD^+^ also serves as a signaling intermediate and only the oxidized form is used as a co-substrate for mitochondrial Sirtuin enzymes, ADP-ribosylases, and NAD^+^ glycohydrolases, many of which have roles in AML (8,9). In these scenarios, the local concentration of NAD^+^ can critically limit enzymatic activities. Additionally, NAD^+^ turnover by these enzymes necessitates import of the molecule into the matrix.

The human SLC25A51 transporter has been identified to selectively import free oxidized NAD^+^ into the mitochondrial matrix (10–12). Loss of SLC25A51 resulted in depleted mitochondrial NAD^+^ levels and its overexpression resulted in elevated concentrations. Thus, SLC25A51 regulates mitochondrial NAD^+^ levels and it is needed for oxidative phosphorylation and mitochondrial ATP production in mammalian cells. Notably, targeting SLC25A51 limited NAD^+^ depletion to the mitochondrial compartment and minimally affected whole-cell or cytoplasmic NAD^+^ levels (10–12). Here, we investigate how mitochondrial NAD^+^ levels contribute to AML pathobiology through modulation of SLC25A51 levels. We hypothesize that the impact of SLC25A51 extends beyond mitochondrial ATP production and simultaneously supports multiple cancer hallmarks including oxidative mitochondrial reactions.

## Results

### Elevated levels of SLC25A51 in AML

We first assessed the relative expression of SLC25A51 from transcriptomic data of AML patient cells (GSE63270 (13)). Compared to hematopoietic stem cells from control healthy adult bone marrow, expression of SLC25A51 was significantly elevated in leukemic Lin-CD34+ CD38+ and Lin-CD34-cells (Fig. 1A). We next evaluated clinical outcomes delineated by expression of SLC25A51. In four distinct databases— Cancer Genome Atlas (TCGA), TARGET AML, as well as AML transcriptome datasets GSE12417 (14) and GSE37642 (15)—cases with the highest SLC25A51 expression (top 25%) were significantly associated with poorer outcomes (Fig. 1B). Broadly, SLC25A51 expression inversely stratified with favorable patient outcomes. To gain insight into SLC25A51’s role in AML cells, we performed gene set enrichment analysis (GSEA) to determine the gene signatures that co-enriched with high SLC25A51 expression in the TCGA datasets. Among the top signatures was the mitochondrially-centered process of oxidative phosphorylation, as defined by its Hallmark gene set (Fig. 1C; Supplementary Fig. S1). Together, the data indicated that import of mitochondrial NAD^+^ by SLC25A51 could be a critical aspect supporting oxidative phosphorylation in AML tumorigenesis. This is consistent with AML’s reliance on oxidative phosphorylation for ATP production (16,17) and SLC25A51’s established role in cellular respiration (10– 12).

**Figure 1.**
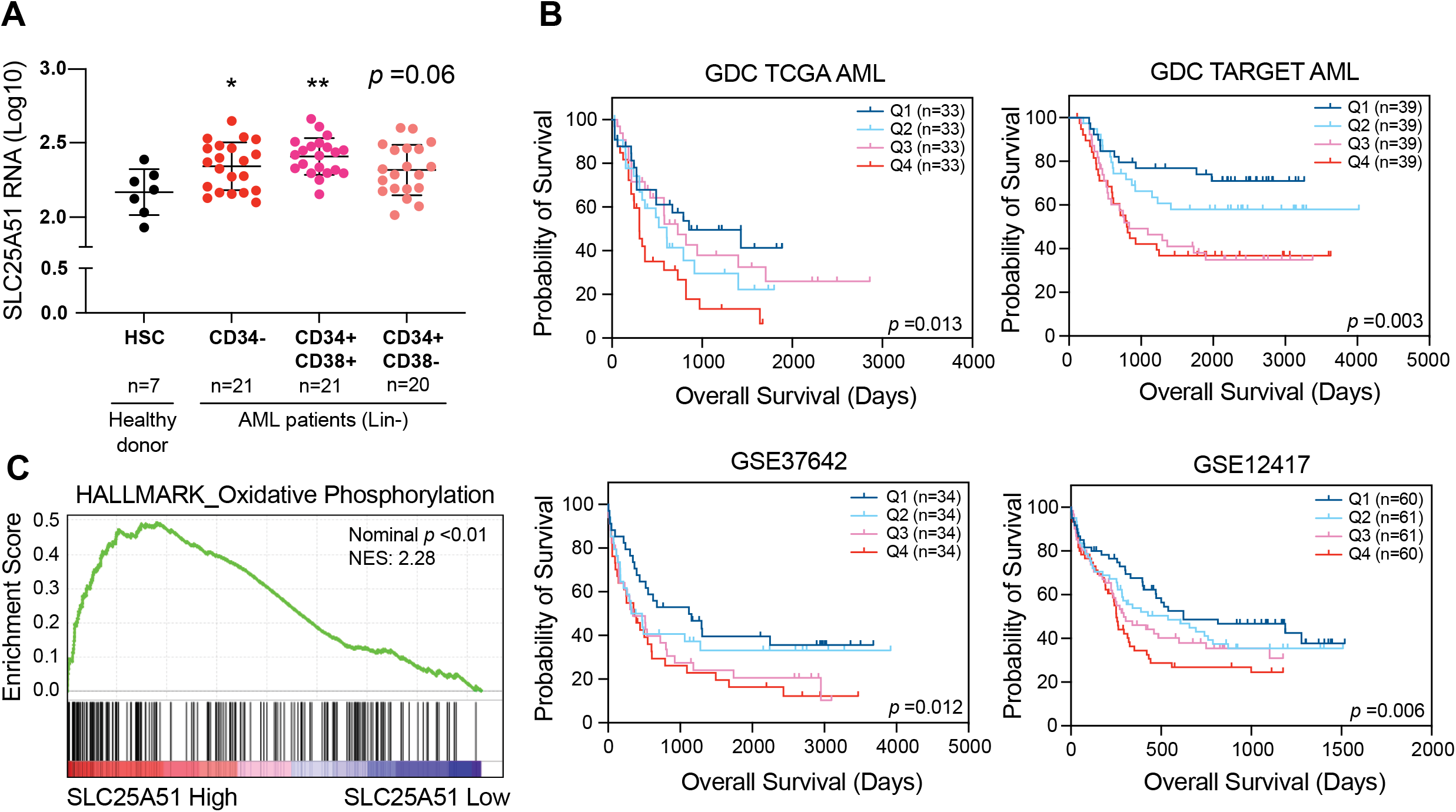
Elevated levels of SLC25A51 in AML. (A) SLC25A51 mRNA expression in hematopoietic stem cell (HSC) from healthy donors and leukemic cells as indicated from AML patients; data from GSE63270. Mean ± SD; ANOVA *p*=0.006, Dunnett’s post-hoc test, \**p* <0.05; \*\**p* <0.01. (B) Kaplan-Meier curves for survival outcomes striated by AML transcriptome datasets (TCGA, TARGET, GSE37642, and GSE12417). Patients were divided into four quadrants according to SLC25A51 mRNA expression. Q1 represents the lowest 25% and Q4 represents the highest 25%. Indicated *p* values were determined by log-rank test. (C) Gene Set Enrichment Analysis using Hallmark gene sets from MSigDB depict the relative enrichment of oxidative phosphorylation associated genes in AML patients with high (top 25%) and low (bottom 25%) SL25A51 expression.

### SLC25A51 is required for AML cell proliferation and survival

To directly test the requirement of SLC25A51 in leukemic cells, we depleted SLC25A51 from a panel of human myeloid cell lines derived from acute (AML, ALL) and chronic (CML, MDS) leukemias using a published short hairpin RNA (shRNA) that had been validated by rescue and Western Blot experiments (10). The panel encompassed lines with a range of genetic aberrancies, childhood and adult leukemias, as well as refractory and relapsed cases (Supplementary Table 1). We observed some variability in expression of SLC25A51 across the lines (Supplementary Fig. S2A and S2B), nevertheless in all cases, its partial depletion resulted in a significant reduction in doubling rate (Fig. 2A; Supplementary Fig. S2C; Supplementary Table S2). We further evaluated clonogenicity and found significantly fewer colonies upon SLC25A51 depletion (Fig. 2B). Notably, MOLM-13 cells were sensitive to partial depletion of SLC25A51, indicating a distinct mechanism compared to Complex I inhibition (2). To gain insight into the diminished fitness of the lines, we performed cell cycle analyses and examined Annexin V dye staining. While there was a statistically significant increase in the percent of cells in G2/M-phase at the expense of S-phase in both shRNA depleted cells and knockout (KO) cells (Supplementary Fig. 2D), the more striking phenotype was increased Annexin V staining at 72 hours post-transduction indicating that cells were directed towards apoptosis (Fig. 2C; Supplementary Fig. S2E). To test whether SLC25A51 was required for AML expansion in vivo, we injected U937 cells depleted for SLC25A51 (1 × 10^6^/mouse) into immunodeficient NOD-*scid* IL2Rg^null^ (NSG) mice. Compared to cells expressing control shRNAs, depletion of SLC25A51 significantly improved median survival (Fig. 2D). Postmortem analyses of bone marrow cells for hCD45^+^ by flow cytometry confirmed that both cohorts had equally robust engraftment (∼40%) (Supplementary Fig. 2F). Thus, the loss of SLC25A51 limited the expansion of AML cells both in vitro and in vivo.

**Figure 2.**
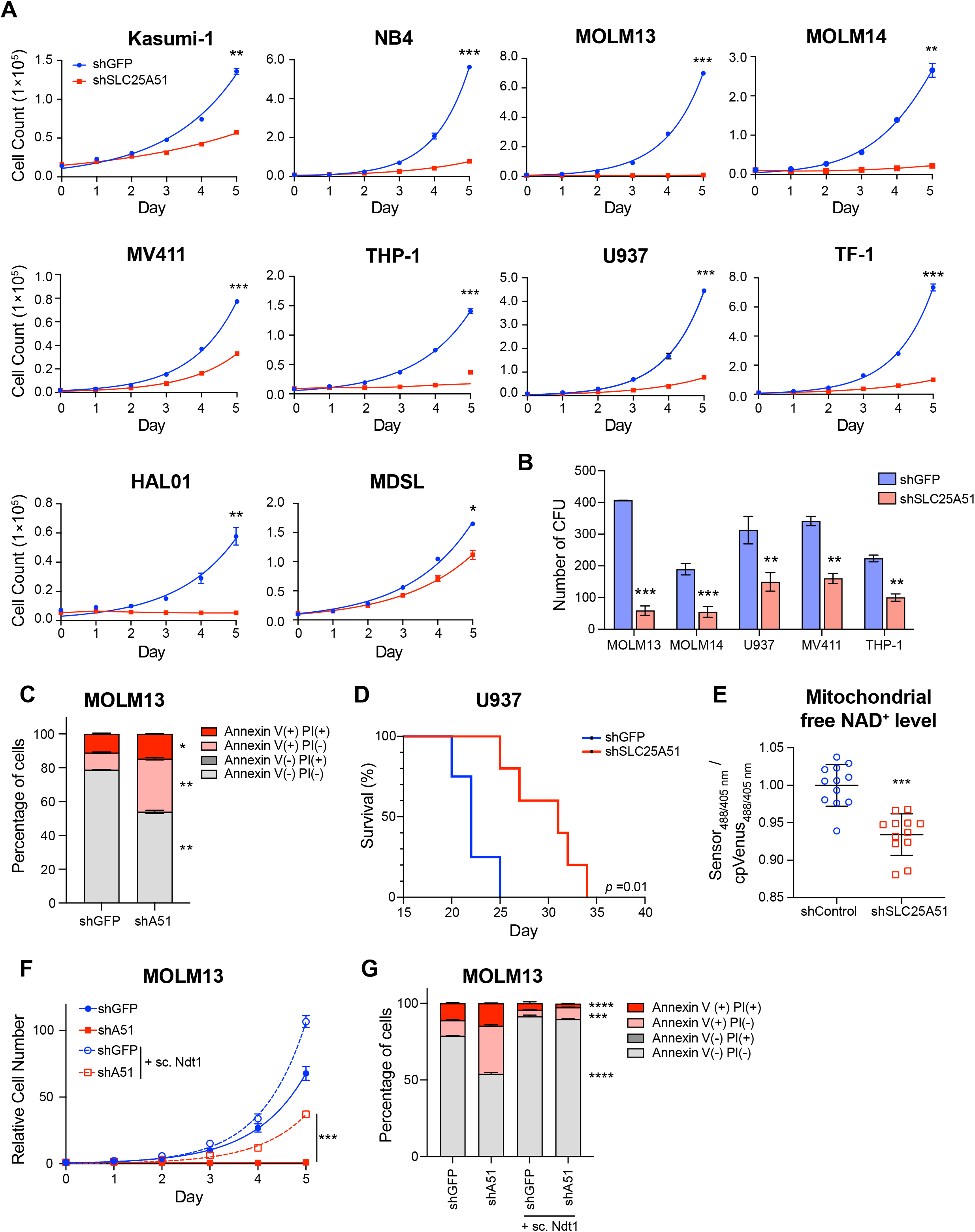
SLC25A51 is required for AML cell proliferation and survival. (A) Growth curves in leukemic cells expressing shSLC25A51 or shGFP (control). Cell counts were measured at indicated times by flow cytometry, mean ± SD (n=3). (B) Clonogenic assays of shSLC25A51 or shGFP expressing AML cells performed in methylcellulose medium. Colony numbers were assessed on day 14 after plating, mean ± SD (n = 3-4). (C) Apoptotic rates in MOLM-13 cells expressing shSLC25A51 or shGFP were measured by flow cytometry using annexin V-FITC and Propidium Iodide (PI) double staining and depicted as relative percentages of viable (annexin V-/PI-), early-stage apoptotic (annexin V+/PI-), late-stage apoptotic (annexin V+/PI+), and dead cells (annexin V-/PI+); mean ± SD (n=3). (D) Kaplan-Meier survival curve following transplantation of 1 × 10^6^ shGFP (n=4) control or shSLC25A51 (n=5) expressing U937 cells into NSG mice; *p*=0.01, log-rank test. (E) Mitochondrial free NAD^+^ levels in MOLM-13 cells expressing shSLC25A51 or shControl scrambled control were measured by mitochondrially-targeted NAD^+^ biosensors (n=12). The fluorescence ratios (488 nm/405 nm) measured by flow cytometry were normalized to cpVenus whose fluorescence is unaffected by NAD^+^ levels. (F) Growth curves of MOLM-13 cells expressing shSLC25A51 or shGFP control in the presence or absence of yeast mitochondrial NAD^+^ transporter, Ndt1; mean ± SD (n=3). (G) Apoptotic rates for shSLC25A51 or shGFP expressing MOLM13 cells measured by annexin V-FITC/Propidium Iodide (PI) double staining and flow cytometry; percentages of viable, early-stage apoptotic, late-stage apoptotic, and dead cells were calculated; mean ± SD (n=3). *p* values in this figure were determined by unpaired, two-tailed Student’s t-test (for two groups) or ANOVA with multiple comparisons analysis using Dunnett’s post-hoc analyses (for groups of three). \**p* <0.05, \*\**p* <0.01, \*\*\**p* <0.001, and \*\*\*\**p* <0.0001 versus control.

### Diminished free mitochondrial NAD^+^ levels are responsible for the observed effects of SLC25A51 loss

To determine whether the observed effects of SLC25A51 depletion were due to loss of mitochondrial NAD^+^, we used the mitochondrially-targeted cpVenus NAD^+^ biosensor (7) in MOLM-13 cells and confirmed that depletion of SLC25A51 diminished mitochondrial free NAD^+^ levels (Fig. 2E). This indicated that AML cells required SLC25A51 activity to actively replenish mitochondrial NAD^+^ for growth of the population. Furthermore, the observed SLC25A51-dependent proliferation defects were rescued by NDT1—a structurally distinct mitochondrial NAD^+^ transporter from yeast (18)—in both SLC25A51-depleted MOLM-13 cells and KO Hap1 cells (Fig. 2F; Supplementary Fig. S2G and S2H). This demonstrated that loss of replenishment was responsible for the phenotypes. Notably, overexpression of NDT1, which elevated steady-state free mitochondrial NAD^+^ levels (10), increased the proliferation of MOLM-13 cells (Fig. 2F; Supplementary Fig. S2G). This indicated that alterations in mitochondrial NAD^+^ levels were sufficient to modulate AML cell proliferation. Ndt1 ectopic expression also rescued apoptotic rates (Fig. 2G). We examined mitochondrial volume and membrane potential and found that in most lines neither was significantly altered by loss of SLC25A51. The exception was the membrane potential in MOLM-13 cells (Supplementary Fig. S2I and S2J).

### Depletion of SLC25A51 targets oxidative mitochondrial processes and alters glutamine utilization

Loss of SLC25A51 blocked ATP production (10–12) similarly to Complex I inhibition but given that its depletion evoked a distinct effect in MOLM-13 cells, we sought to determine the major pathways impacted by loss of mitochondrial NAD^+^.

We observed elevated mitochondrial superoxide levels upon depletion of SLC25A51 (Fig. 3A) and considered that an altered redox state could create inefficient or reversed flux through mitochondrial dehydrogenase enzymes and ETC complexes. To determine the oxidation ratio of SLC25A51-depleted mitochondria, we obtained simultaneous measurements of coenzyme Q_10_ (CoQ_10_) states. CoQ_10_ strictly localizes to mitochondria and the relative abundances of ubiquinone (oxidized) and ubiquinol (reduced) indicate the local oxidoreductive environment (19). We observed specific loss of ubiquinol species without significant change in total CoQ_10_ levels (Fig. 3B). This indicated a lowered oxidation ratio and a disrupted flow of the electron transport chain, both consistent with the observed loss of oxidative respiration.

**Figure 3.**
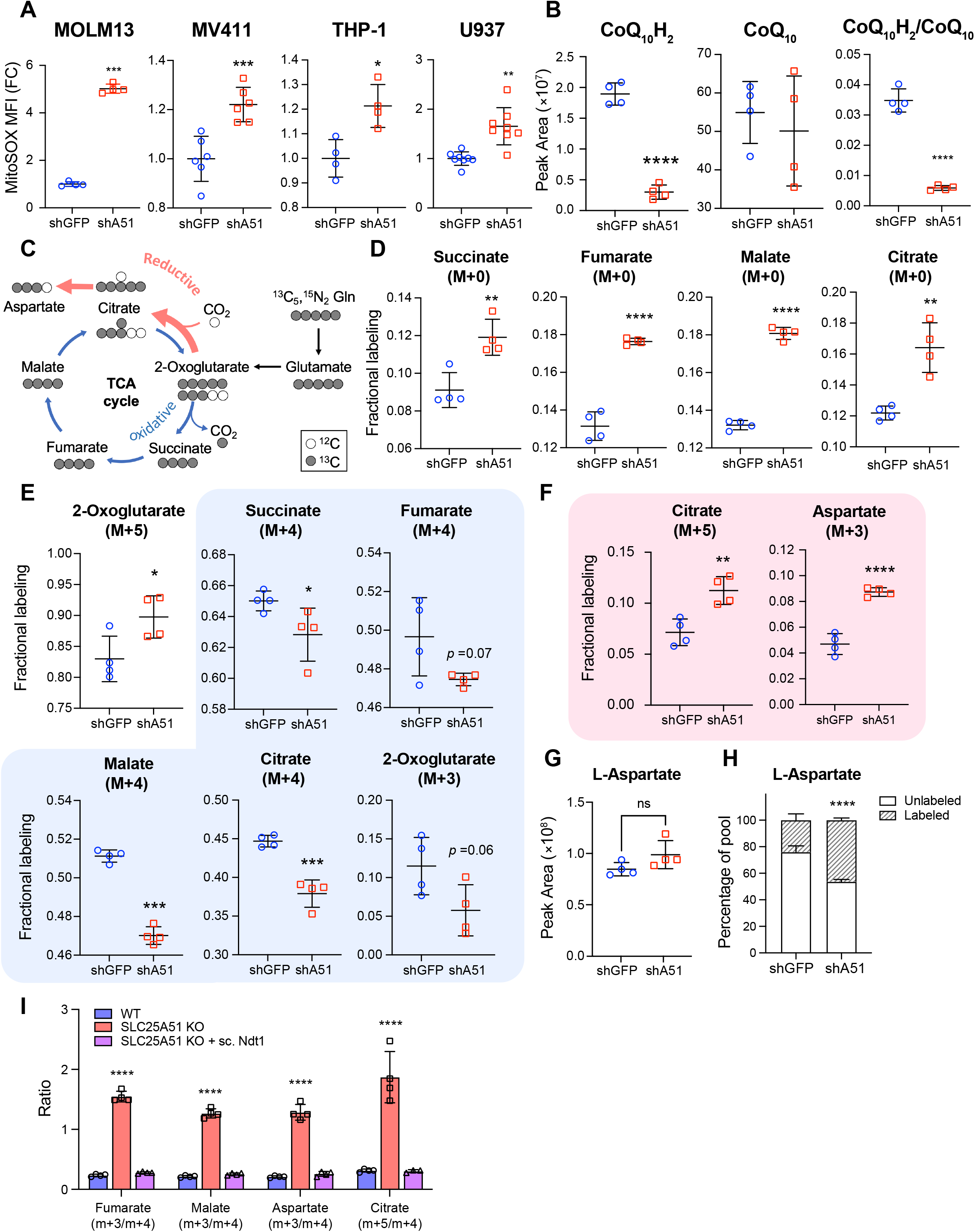
Depletion of SLC25A51 targets oxidative mitochondrial processes. (A) Mitochondrial superoxide levels were measured in shSLC25A51 or shGFP expressing AML cells using MitoSOX™ Red staining and flow cytometry; mean ± SD (n=4-8). (B) CoQ_10_H_2_/CoQ_10_ ratio and total levels were measured in shSLC25A51 or shGFP expressing U937 cells using UHPLC-MS/MS; mean ± SD (n=4). (C) Schematic depiction of the oxidative (blue) and reductive (red) TCA metabolites derived from [^13^C_5_,^15^N_2_]-glutamine tracing experiments. (D) Stable isotope-resolved metabolomics analysis performed in U937 cells expressing either shSLC25A51 and shGFP and equilibrated with media-containing [^13^C_5_,^15^N_2_]-glutamine for 24h. Fractional labeling of M+0 succinate, fumarate, malate, and citrate; mean ± SD (n=4). (E) Fractional labeling of 2-oxoglutarate (M+5) and oxidative TCA metabolites including succinate (M+4), fumarate (M+4), malate (M+4), citrate (M+4), and 2-oxoglutarate (M+3); mean ± SD (n=4). (F) Fractional labeling of reductive TCA metabolites including citrate (M+5) and aspartate (M+3) derived from [^13^C_5_,^15^N_2_]-glutamine; mean ± SD (n=4). (G) The total levels of L-aspartate in shSLC25A51 and shGFP expressing U937 cells was measured by UHPLC-MS/MS; mean ± SD (n=4). (H) The percentage of aspartate derived from [^13^C_5_,^15^N_2_]-glutamine in shSLC25A51 and shGFP control U937 cells were measured by UHPLC-MS/MS; mean ± SD (n=4). (I) The ratio of oxidative to reductive TCA metabolites including fumarate, malate, aspartate, and citrate derived from [^13^C_5_,^15^N_2_]-glutamine; mean ± SD (n=4). *p* values in this figure were determined by unpaired, two-tailed Student’s t-test (for two groups) or ANOVA with post hoc Dunnett’s multiple comparisons analysis (for groups of three). \**p* <0.05, \*\**p* <0.01, \*\*\**p* <0.001, and \*\*\*\**p* <0.0001 versus control.

AML depends on glutaminolysis (3,20,21), so we labeled proliferating U937 cells with [U-^13^C, U-^15^N]-glutamine and analyzed its relative incorporation at 24 hours in equilibrated cells that were depleted for SLC25A51 compared to control (Fig. 3C). In U937 cells, we found that high levels of SLC25A51 was required for robust glutaminolysis and anaplerosis. Depletion of SLC25A51 resulted in increased usage of non-glutamine carbon sources to support the TCA cycle, as measured by increased proportions of unlabeled TCA intermediates (Fig. 3D). Furthermore, SLC25A51 was required to sustain oxidative TCA flux (Fig. 3E). This was observed through decreased levels of M+4 labeled TCA intermediates and M+3-labeled oxoglutarate. In contrast, levels of M+5 labeled citrate increased, indicating an increase in reductive flux. This change in flux was especially clear with M+3 aspartate (Fig. 3F), a critical metabolic product for cycling cells (22–24). While there was no significant change in overall aspartate levels, we observed an increased fraction derived from reductive (M+3) glutaminolysis (Fig. 3G and 3H). The observation of elevated reductive TCA intermediates following loss of SLC25A51 activity was recapitulated in CML-derived Hap1 SLC25A51-knockout (Hap1-KO) cells (10) and rescued by expression of Ndt1 (Fig 3I; Supplementary Fig. S3A). Hap1-KO cells further had a significant increased proportion of aspartate derived from reductive reactions (Supplementary Fig. S3A).

We reasoned that because import of mitochondrial NAD^+^ could restore proliferation and oxidative TCA cycle flux (Fig. 2F, Fig 3I; Supplementary Fig. S2F, S2G, and S3A) it may do so by controlling the NAD^+^/NADH ratio—rather than absolute NAD^+^ concentrations. We expressed ^Mito^*Lb*Nox (25) to test whether oxidizing the remaining mitochondrial NADH to NAD^+^ impacted AML cells exhibiting deteriorating fitness with depleted levels of SLC25A51. This approach was feasible with the shRNA because mitochondrial NAD levels were only partially depleted. We found that increasing the oxidized mitochondrial NAD^+^/NADH ratio restored proliferation in AML cells (Supplementary Fig. S3B). Together, this indicated that changes in SLC25A51 activity can control directional flux of the TCA cycle by controlling the mitochondrial NAD^+^/NADH ratio and its oxidative state. Thus, SLC25A51 is a regulatory point for oxidative TCA metabolism in AML.

### Minimized impact on pathways outside of mitochondria

Although SLC25A51-depletion decreased oxidative flux and promoted reductive TCA reactions, we found that a level of SLC25A51-depletion capable of impacting oxidative flux did not significantly deplete total aspartate levels and did not deplete AMP or GMP levels (Supplementary Fig. S3C). Furthermore, there were minimal effects on levels of glycolytic intermediates, including lactate levels, as well as 2-hydroxyglutarate (Supplementary Fig. S3D).

### Depletion of SLC25A51 sensitizes AML cells to glutamine deprivation

We had observed that depletion of SLC25A51 forced AML cells to more heavily utilize glutamine for the synthesis of aspartate, at the expense of oxidative TCA reactions. We therefore reasoned that loss of SLC25A51 could be combined with glutamine deprivation as an approach to dampen proliferation in cells that characteristically depend on oxidative glutaminolysis. We first tested whether depletion of SLC25A51 sensitized cells to removal of glutamine from the growth media. We found that its loss significantly impaired proliferation of AML upon depletion of SLC25A51 compared to control (Fig. 4A; Supplementary Fig. S4A; Supplementary Table S3). In agreement, SLC25A51 depletion sensitized cells to 1µM CB-839 in growth curve measurement for 5 days, a selective inhibitor of glutaminase enzymes used to deaminate glutamine for intracellular use (Fig. 4B; Supplementary Fig. S4B; Supplementary Table S4). We further observe increased apoptotic percentages when SLC25A51 depletion was combined with CB-839 treatment (Fig. 4C; Supplementary Fig. S4C).

**Figure 4.**
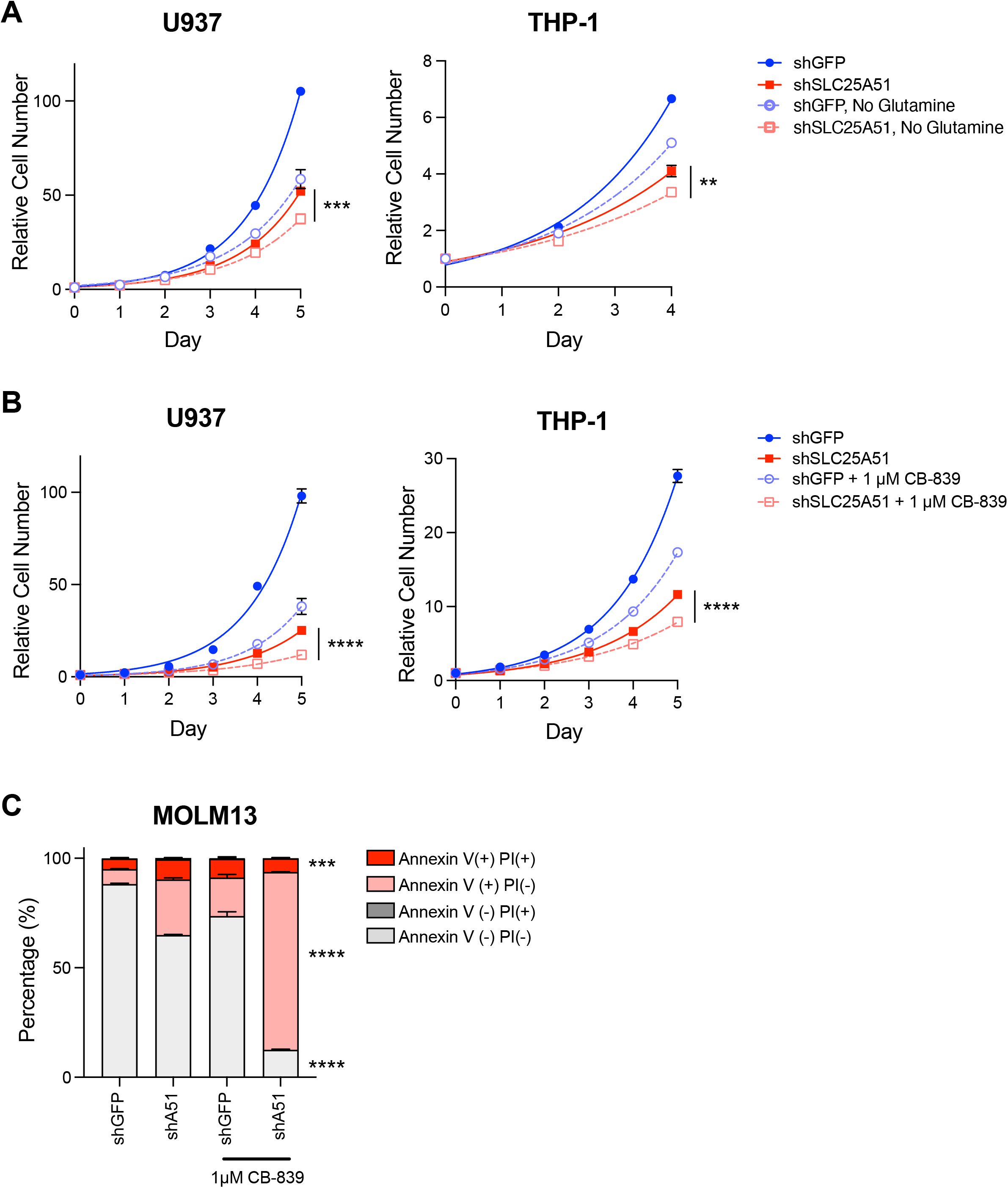
Depletion of SLC25A51 sensitizes AML cells to glutamine deprivation. (A) Growth curves of shSLC25A51 or shGFP expressing AML cells grown with indicated concentrations of glutamine. Cell counts were measured at indicated times by flow cytometry; mean ± SD (n=3). (B) Growth curves of shSLC25A51 or shGFP expressing AML cells treated with 1µM CB-839 for 5 days. Cell counts were measured as above; mean ± SD (n=3). (C) Apoptotic rates of shSLC25A51 or shGFP expressing MOLM-13 cells treated with 1µM CB-839 and analyzed by annexin V-FITC/Propidium Iodide (PI) staining using flow cytometry on day 5; mean ± SD (n=3). *p* values in this figure were determined by unpaired, two-tailed Student’s t-test at end point. \*\**p* <0.01, \*\*\**p* <0.001, and \*\*\*\**p* <0.0001 versus shSLC25A51 group.

## Discussion

NAD^+^ sensors (7,26) have demonstrated that the human SLC25A51 transporter was required to sustain free mitochondrial NAD^+^ levels (10), and thus depletion of SLC25A51 provided a unique ability to selectively study the tumorigenic contribution for replenishment of this dinucleotide in mitochondria. Here, we showed that SLC25A51 can serve as a regulatory point in AML by controlling import of oxidized mitochondrial NAD^+^. Expression of SLC25A51 was upregulated in leukemic cells and its higher expression correlated with poorer outcomes in AML patients (Fig. 1A and 1B). Metabolic flux analyses showed that higher SLC25A51 levels promoted glutamine-fueled oxidative flux through the TCA cycle (Fig. 3E and F, 3I), and thereby provided a mechanism for how diminished SLC25A51 levels impaired AML expansion in vitro and in vivo. This was further supported by rescued proliferation and viability of AML cells from the ectopic expression of either Ndt1 to restore free mitochondrial NAD^+^ levels or ^Mito^*Lb*Nox to favor an oxidative NAD^+^/NADH ratio in mitochondria (Supplementary Fig. 3B).

In this scenario, an oxidative NAD^+^/NADH ratio was critical for robust proliferation. Nevertheless, depletion of SLC25A51 is expected to simultaneously affect multiple pathways. In addition to controlling oxidative TCA flux, loss of SLC25A51 would impair other oxidoreductive reactions in mitochondria such as fatty acid oxidation (27), which is a robust source of carbon for many tumors (4). Moreover, the activities of resident NAD^+^ consuming enzymes—e.g. mitochondrial SIRT3, SIRT4, and SIRT5— would directly depend on the NAD^+^ imported by SLC25A51 (6). Compellingly, drug-resistant AML has been found to depend on fatty acid oxidation and SIRT5 (28,29). Thus, simultaneously impacting multiple pathways with a single target has potential for reducing the chances of adaptation, resistance, and relapse of the disease. It may also be applicable for broadly sensitizing AML that arise from distinct mechanisms.

The results of this study are in line with a growing appreciation that many cancers rely on upregulation of NAD^+^ biosynthetic pathways (30–32) and mitochondrial activity for growth advantages. In the case of leukemias, there are already several chemotherapeutics with targets related to mitochondrial function, including Venetoclax (33) targeting BCL-2 proteins and multiple electron transport chain Complex I inhibitors, including IACS-010759 (2) and Mubritinib (34). Interestingly, loss of SLC25A51 desensitized AML cells to Mubritinib treatment (34).

Whole-cell loss of NAD^+^ through nicotinamide phosphoribosyltransferase inhibition was shown to be a potent vulnerability for drug-resistant AML (35,36) but this approach is broadly considered as likely having too narrow of a therapeutic window to be implemented in practice. In contrast, loss of SLC25A51 resulted in selective depletion of mitochondrial NAD^+^ levels and minimally affected whole cell levels. Thus, targeting SLC25A51 could represent an approach to precisely impair AML cells both via targeting their heightened dependence on oxidative NAD^+^ as well as by limiting its depletion to the mitochondrial organelle.

In support, we have shown that there exists a window for SLC25A51 partial depletion that effectively impairs the expansion of AML cells but had minimal effects on total AMP or GMP levels and levels of glycolytic intermediates, including lactate (Supplementary Fig. 3C and 3D). It is thus possible that SLC25A51 targeting may be relatively well tolerated compared to whole cell NAD^+^ depletion and be able to avoid some of the observed toxicity with Complex I inhibitor IACS-010759 such as nucleotide depletion and lactic acidosis (2).

We surmise that AML cells may be exquisitely sensitive to loss of SLC25A51 because of its impact on multiple oxidative pathways that would otherwise compensate for each other. Moreover, the resulting compartmentalized depletion of mitochondrial NAD^+^ appeared to support at least some continued nucleotide and amino acid synthesis (Fig. 3E and F). What was altered, nevertheless, was the source of carbon used to support amino acid and nucleotide synthesis (Fig. 3D); the carbon source for lipid synthesis was undetermined.

In this regard, depletion of SLC25A51 is mechanistically distinct from Complex I inhibition. SLC25A51 depletion in the U937 AML cell line resulted in increased incorporation of non-glutamine carbon into TCA intermediates compared to Complex I block by IACS-010759 (2), indicating that high SLC25A51 in AML promoted glutamine incorporation into the TCA cycle. We also observed that in vitro proliferation of AML cells with high glycolytic capacity such as MOLM-13 were distinctly sensitive to SLC25A51 depletion. This may be due to incorporation of glucose-derived carbons, in lieu of glutamine, into TCA intermediates but this still needs to be determined. Interestingly, in CML-derived Hap1 cells, altered glutamine usage was not readily observed from loss of SLC25A51 (Supplementary Fig. S3A), underscoring metabolic differences between different leukemia lines (37).

When SLC25A51 levels were lowered in AML cells, we observed increased incorporation of glutamine carbons into aspartate. There was minimal change in total aspartate levels, suggesting that depletion of SLC25A51 shunted glutamine flux away from supporting the TCA cycle and towards support of aspartate synthesis required for proliferation (22–24) (Fig. 3G and H). Thus, we reasoned that SLC25A51 depletion could sensitize AML cells to glutamine deprivation, and this was the rationale for testing whether SLC25A51 loss could potently combine with CB-839 treatment (Fig. 4B, Supplementary Fig. 4B and Supplementary Table 4). The data suggested that partial SLC25A51 depletion may be an effective approach to selectively sensitize AML cells to current therapies and further help with treatment tolerance.

Given that the data indicated excess mitochondrial NAD^+^ may serve a pathological role in AML, it will be critical to further investigate the full impact of NAD^+^ supplementation that is currently being investigated as an adjuvant to chemotherapy and to support cell health during immunotherapy (38–41). While this study directly explored SLC25A51 in AML cells, our findings provide a rationale for more broadly exploring its roles in other tumor types that depend on oxidative mitochondrial metabolism. A recent study identified that an oxidative mitochondrial NAD^+^/NADH ratio was critical in ovarian cancer (42), and another reported that high SLC25A51 levels in hepatocellular carcinoma (HCC) promoted cancer progression through SIRT5 activation (43). Additionally, there are some phenotypic similarities between SLC25A51 loss in AML and block of mitochondrial pyruvate carrier in oxidative Non-Hodgkin Diffuse Large B Cell Lymphomas (44). It is possible that these carriers may function together to promote oxidative TCA flux in this blood cancer as well.

In conclusion, our data indicated that SLC25A51 supports AML expansion by promoting glutamine incorporation sustaining an oxidative TCA cycle. We have provided a rationale for future work to determine effectiveness and toxicity of targeting SLC25A51 for AML where there is an unmet need for better tolerated therapeutic options, especially with elderly patients.

## Author contributions

M-J.L. and X.A.C. conceived and designed the overall study. All authors (M-J.L., J.B., V.I, Y-T.C., W.M., S.T., and X.A.C) contributed to the development of experimental approaches, analyzed data, and interpreted the experiments. M-J.L., J.B., V.I., and Y-T.C performed experiments. M-J.L. and X.A.C wrote the manuscript. All authors reviewed and edited the manuscript.

## Acknowledgements

This work was supported by grants from the Cancer Prevention and Research Institute of Texas (RP210079), the PEW Charitable Trusts and National Institutes of Health (DP2GM126897 and R01 CA206210).

## Data Availability Statement

The original data generated in this study are available within the article, its supplementary data files, or upon request from the corresponding author. Expression profile data analyzed in this study were obtained from TGCA, TARGET, and Gene Expression Omnibus (GEO) at GSE63270, GSE37642, and GSE12417.

## Materials and Methods

### Mice

All animals were treated and maintained in the University of Texas at Austin animal resources center. All animal procedures were performed according to the guidelines approved by the Institutional Animal Care and Use Committee (IACUC) at University of Texas at Austin, protocol AUP-2020-00249.

### Cell culture

NB4, MOLM-13, MOLM-14, MV-411, THP-1, U937, and HAL01 cells were cultured in RPMI 1640 medium containing 2 mM L-glutamine, supplemented with 10% fetal bovine serum and 1× penicillin-streptomycin. Kasumi-1 cells were cultured in RPMI 1640 medium containing 2 mM L-glutamine, supplemented with 20% fetal bovine serum and 1× penicillin-streptomycin. TF-1 and MDS-L cells were cultured in RPMI 1640 medium containing 2 mM L-glutamine, supplemented with 10% fetal bovine serum, 1× penicillin-streptomycin, and 2ng/mL GM-CSF and 10ng/mL hIL-3 respectively. HAP1 wild-type (Horizon Discovery: C631) and HAP1 SLC25A51-KO 4-bp deletion (Horizon Discovery: HZGHC001927c010) cells were cultured in Iscove’s modified Dulbecco’s culture medium supplemented with 10% fetal bovine serum and 1× penicillin–streptomycin.

### Lentivirus production

Parental HEK293T cells were cultured in Dulbecco’s modified Eagle’s medium (DMEM) containing 4.5 g/L glucose, 1 mM sodium pyruvate and 4mM L-glutamine, supplemented with 10% fetal bovine serum and 1× penicillin-streptomycin. Lentiviruses were generated by co-transfection of shRNA constructs with psPAX2 (packaging) and pMD2.G (envelope) into HEK293T cells. Supernatants were collected 60 h after transfection, filtered through 0.45um PVDF syringe filter, and concentrated by using polyethylene glycol (PEG) 6000. Briefly, 42.5% of PEG 6000 solution was prepared in sterile water with 1.5M NaCl, mixed in a 1:4 ratio with the virus-containing supernatant and incubated for 4 hours at 4°C. Subsequently, the mix was spun down for 30 minutes at 6,000 x g at 4°C. The pellets were then resuspended in a suitable volume of sterile PBS with 1% BSA, aliquoted, and stored at −80 °C.

### shRNA knockdown

Cells were transduced with lentivirus encoding shRNA targeting against human SLC25A51 (Sigma, Mission shRNA). shRNA sequences used are: TRCN0000060235, CCGGGCAACTTATGAGTTCTTGTTACTCGAGTAACAAGAACTCATAAGTTGCTTTTTG. Control shGFP sequence: GCAAGCTGACCCTGAAGTTCAT. Control non-targeting scramble sequence: CCTAAGGTTAAGTCGCCCTCG. To generate stable knockdown cell lines, cells were transduced with lentiviruses encoding shRNA and selected with 1-2 µg/mL puromycin.

### Cell proliferation rate calculation

Cells were counted and plated in triplicate in 12-well plates, with initial seeding density of 10,000 cells per well. Viable cells were counted daily using a NovoCyte flow cytometer and proliferation rate was calculated using following formula (24): Proliferation Rate (Doublings per day) = log_2_ (Final cell count/Initial cell count)/ days

### Methylcellulose culture

Cells (1,000 cells/per 35 mm dish) were plated and cultured in methylcellulose medium (MethoCult H4230) for 14 days. Colonies containing of more than 40 cells were counted as a colony using microscopy.

### Apoptosis measurement

Apoptotic rate was measured by using FITC Annexin V Apoptosis Detection Kit I (BD556547). Briefly, cells were counted and resuspended in 1× Annexin V binding buffer at a concentration of 1×10^6^ cells/ml. 100 µl (1×10^5^ cells) of cell suspension was stained with FITC Annexin V and PI at RT for 15 minutes in the dark. Subsequently, 400 µl of 1× Annexin V binding buffer was added to each tube and analyzed by NovoCyte flow cytometer within an hour.

### Cell cycle analysis

Cells were fixed with ice-cold 75% ethanol at −20 °C overnight. Following two washes with staining buffer (PBS with 1%FBS), the cells were stained with Ki-67 and propidium iodidie (PI) at room temperature (RT) for 30 minutes in the dark and analyzed with a NovoCyte flow cytometer. The cell cycle phase distribution was calculated using FlowJo v.10.

### Real-time quantitative RT-PCR

RNA was isolated with the RNeasy plus mini kit (QIAGEN) following manufacturer’s protocol. cDNA was synthesized using the iScript reverse transcription supermix kit (Bio-Rad #1708841). Quantitative real-time PCR was performed with the Applied Biosystems ViiA 7 system using iTaq Universal SYBR Green Supermix (Bio-rad #1725121). Human actin was used as a house keeping control. To calculate fold change in mRNA expression, the 2^−ΔΔ*C*t^ method was used. Primers were: endogenous human *SLC25A51* mRNA, forward, CGCTGATGGGAAATCCAGTTA, reverse, CTGGAGTTTGGCAGGATGATAG; human β*-actin*, forward, TCCCTGGAGAAGAGCTACG, reverse, GTAGTTTCGTGGATGCCACA.

### Immunoblot analysis

Cells were washed with ice-cold PBS and lysed directly in 2× Laemmli sample buffer containing DTT. Protein samples were electrophoresed on 10% Bis-Tris gels (Invitrogen) and transferred to 0.45 µm nitrocellulose membrane. Membranes were blocked with 5% BSA in Tris-buffered saline (TBS) pH 7.6 containing 0.1% Tween-20 (TBST). Antibodies were prepared in 1% BSA in TBST. Dilutions were as follows: anti-SLC25A51 (MyBiosource, MBS1496255, 1:1,000), anti-α-tubulin (Sigma, T9026, 1:3,000), anti-rabbit IgG H&L IRDye 800CW (Abcam, ab216773, 1:10,000), and anti-mouse IgG H&L Alexa Fluor 680 (Invitrogen, A10038, 1:10,000). Membranes were imaged using the LiCOR Odyssey CLx.

### Mitochondrial NAD^+^ measurement using the NAD^+^ biosensor

MOLM-13 cell line stably expressing ^mito^cpVenus or ^mito^Sensor were transduced with scramble or human SLC25A51 shRNA for 72h. Measurements of mitochondrial NAD^+^ levels using the NAD^+^ biosensor has previously been described (7). In brief, MOLM13 ^mito^cpVenus or ^mito^Sensor cells were collected in cold RPMI and kept on ice until analysis. Data was collected on a NovoCyte flow cytometer using following parameters: excitation 488 nm, emission 530 ± 30 nm and excitation 405 nm, emission 530 ± 30 nm. Cells were gated exclude debris and doublet and 10,000 fluorescent cells were analyzed per sample. Ratiometric 488/405 nm fluorescence values were obtained for each cell using the derived function of FlowJo v.10.

### U937 cell line xenografts

NOD.Cg-Prkdc^SCID^Il2rg^tm1Wjl^/SzJ (NSG) mice (8 weeks old, male, Charles River) were injected intravenously with 1 × 10^6^ U937 cells transduced with shRNA targeting to GFP (as control) or SLC25A51 via tail vein using 29 1/2-gauge needles and followed up for tumor burden. In brief, 1 × 10^6^ cells were suspended in 100uL DPBS and injected into the tail vein of each mouse on day 0. Mice were considered moribund and euthanized in case of rapidly weight loss (greater than 20%, measured daily) and other signs of poor health, including hind limb paralysis or hunched posture. Bone marrow cells were collected and analyzed for human CD45-Pacific blue on NovoCyte flow cytometer.

### Survival analysis

AML patients from TCGA, TARGET, GSE37642, and GSE12417 dataset has corresponding SLC25A51 expression levels were used for analysis. Among these patients, quartiles of expression values were calculated, and overall survival (OS) was estimated using Kaplan-Meier method and compared across these four groups using a log-rank test in each dataset.

### Metabolic flux analysis

Cells were cultured in RPMI-1640 media supplemented with 10% dialyzed FBS, 2 mM [^13^C_5_,^15^N_2_]-glutamine (Cambridge Laboratories) and 4.5 g/L unlabeled glucose. After 24 h, cells were centrifuged, washed twice in cold PBS, and flash-frozen in liquid nitrogen for mass spectrometry analysis as previously described (19,45). Polar and apolar metabolites were extracted using a modified Bligh-Dyer procedure and analyzed by an ultra-high pressure liquid chromatography Vanquish tandem Q-Exactive mass spectrometer system (Thermo Scientific, Waltham, MA). Polar samples were first analyzed using a SeQuant ZIC-HILIC 3.5 μm, 100 Å, 150 × 2.1 mm PEEK coated HPLC column (Millipore Sigma, Burlington, MA) and then analyzed using a Synergi 4 µm Hydro-RP 80 Å, 150 × 2 mm HPLC column (Phenomenex, Torrance, CA) as recently reported (PMID: 32027051, PMID: 34518298). Apolar samples were dried and resuspended in 95:5 ethanol:6N HCl and then analyzed using a Synergi 4 µm Hydro-RP 80 Å, 150 × 2 mm HPLC column (Phenomenex, Torrance, CA) with isobaric separation (PMID: 29475487). Raw MS data was processed using Sieve 2.2 (Thermo Fisher Scientific) and cleaned according to a 25% coefficient of variance threshold. Peaks were then scaled according to probabilistic quotient normalization and features were mined by matching accurate masses and retention times to a library of standards generated in house that include IROA 300 MS Metabolite Library of Standards (IROA Technologies). In addition, accurate masses taken from the Kyoto Encyclopedia of Genes and Genomes (46) and the Human Metabolome databases (47) were also mined.

### Statistical analysis

For statistical analyses, *p* values were determined by applying the two-tailed Student’s t test (for comparison of two groups), one-way ANOVA (for comparison of three or more groups), or log-rank test (for survival curve). All results are presented as mean ± SD as indicated and \**p* <0.05, \*\**p* <0.01, \*\*\**p* <0.001, and \*\*\*\**p* <0.0001. All analyses were performed with GraphPad Prism 9.0 software.

## SUPPLEMENTARY DATA FOR

**Figure S1.**
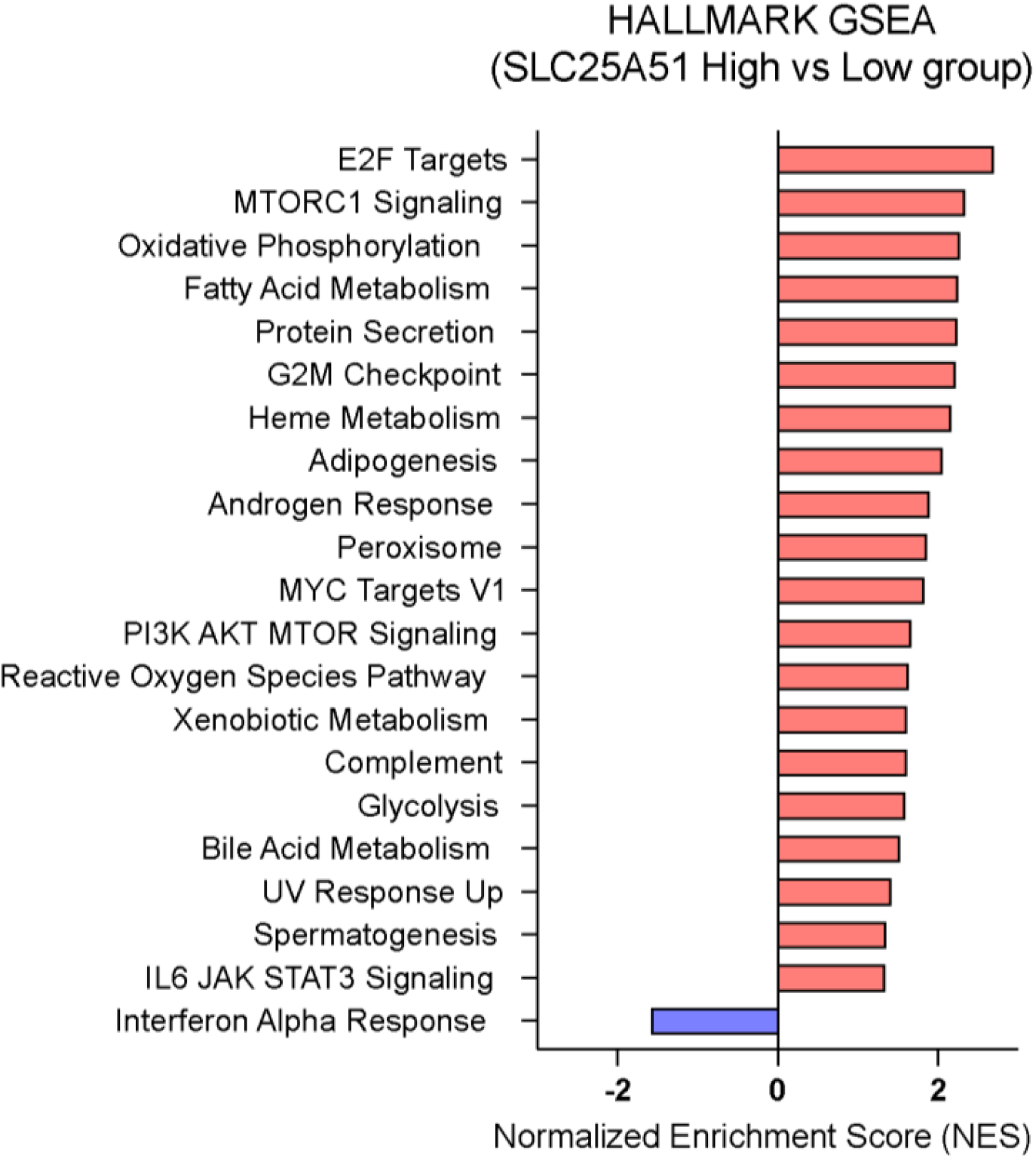
Gene set enrichment analysis (GSEA) results for SLC25A51-high and SLC25A51-low AML patients. Using Hallmark gene sets from the Molecular Signature Database, statistically significant gene sets were selected (FDR < 0.25 and nominal *p*-value <0.05) and ordered by normalized enrichment score (NES). A positive normalized enrichment score (NES) value indicates enrichment in the SLC25A51 high AML patients.

**Table S1.** Panel of human myeloid leukemia cell lines.

| **Name** | **Disease types** | **Sex** | **FAB** |  | **Fusion gene** |
| --- | --- | --- | --- | --- | --- |
| MOLM-13 | AML | M | M5 | Relapse | MLL-AF9 w/FLT3-ITD |
| MOLM-14 | AML | M | M5 | Relapse | MLL-AF9 w/FLT3-ITD |
| U937 | AML | M | M5 | Refractory | CALM-AF10 |
| MV4-11* | AML | M | M5 | Diagnosis | MLL-AF4 w/FLT3-ITD |
| THP-1* | AML | M | M5 | Relapse | MLL-AF9; CSNK2A1-DDX39B |
| NB4 | AML | F | M3 | Relapse | PML-RARA |
| Kasumi-1* | AML | M | M2 | Relapse | AML1-ETO |
| HAL-01 | ALL | F |  |  | E2A-HLF |
| MDSL | Myelodysplastic syndrome | M |  |  |  |
| TF-1 | Acute erythroid leukemia | M | M6 |  | CBFA2T3-ABHD12 |

Abbreviations: F, female; M, male; \* childhood

**Figure S2.**
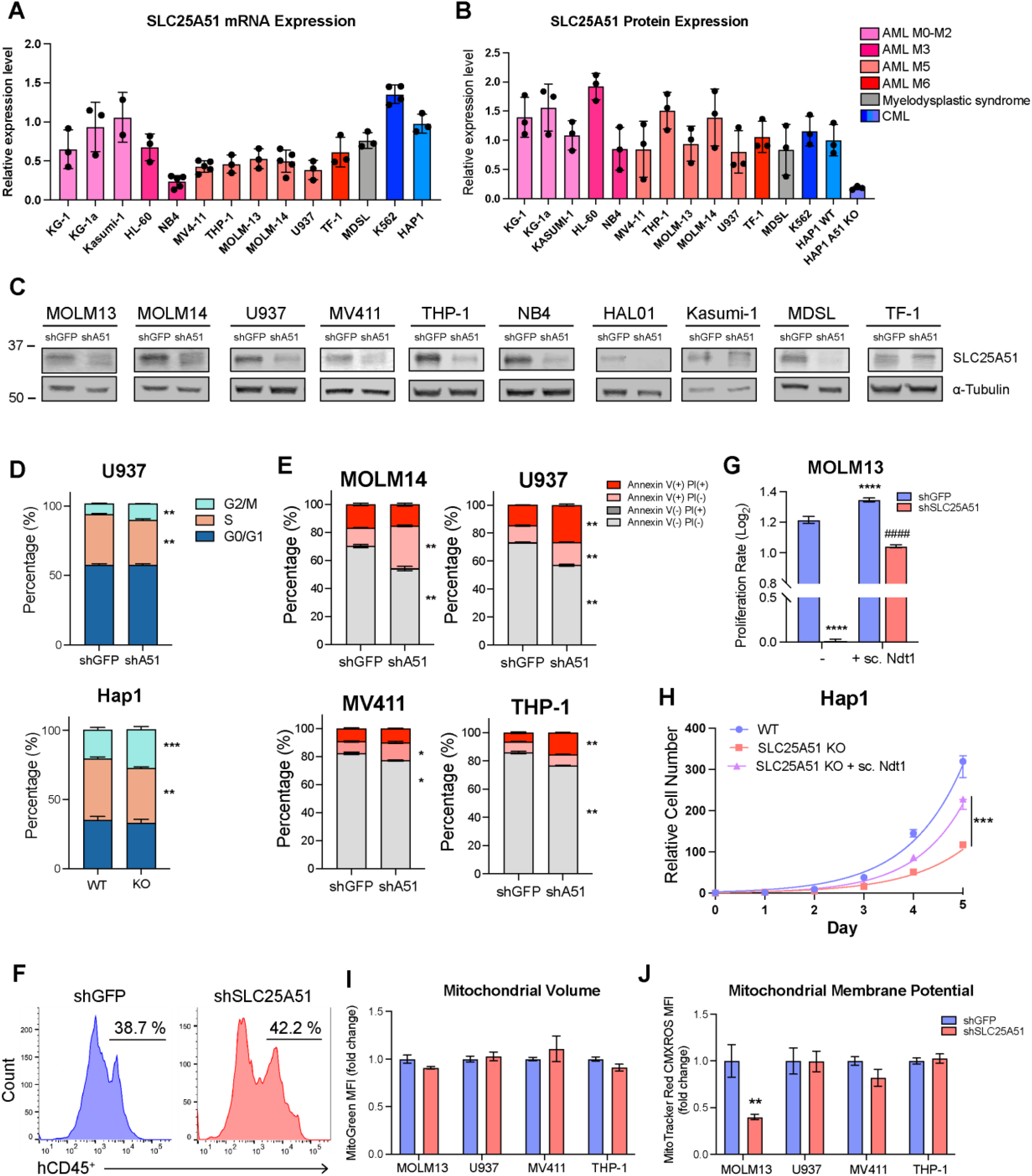
Depletion of SLC25A51 affects proliferation and survival due to the diminishment of mitochondrial NAD^+^ level. (A) SLC25A51 mRNA expression levels were determined by qPCR in human myeloid leukemia cell lines. Relative expression levels were normalized to HAP1 cell line. Data shown as mean ± SD (n=3). (B) SLC25A51 protein expression levels were determined by western blot in human myeloid leukemia cell lines. Relative expression levels were normalized to Hap1 WT cell line. Data shown as mean ± SD (n=3). (C) Knock down efficiency of shSLC25A51 or shGFP expressing AML cells analyzed by Western Blot. (D) Cell cycle analyses were performed using Ki-67 and Propidium Iodide (PI) staining in U937 and Hap1 cells and analyzed by flow cytometry using univariate cell cycle model. Data shown as mean ± SD (n=3). (E) Apoptotic rates in shSLC25A51 or shGFP expressing AML cells were measured by flow cytometry using annexin V-FITC and Propidium Iodide (PI) staining. The percentage of living, early-stage apoptotic, late-stage apoptotic, and dead cells were calculated. Data shown as mean ± SD (n=3). (F) Postmortem analyses of bone marrow collected and analyzed by flow cytometry for the percentage of hCD45^+^ cells. Representative histogram plots shown for both groups. (G) Calculated proliferation rates of shSLC25A51 or shGFP expressing MOLM-13 cells with ectopic expression of yeast Ndt1 transporter. Cell counts on day 5 were normalized to cell numbers on day 0 and used to calculate proliferation rate as described in methods. Data shown as mean ± SD (n=3). (H) Growth curve of Hap1 WT, SLC25A51 KO, and SLC25A51 KO + Ndt1 cells. Cell counts were measured at indicated times by flow cytometry. Data shown as mean ± SD (n=3). (I) Relative mitochondrial volume was measured in shSLC25A51 or shGFP expressing AML cells by flow cytometry using MitoGreen staining. Data shown as mean ± SD (n=3-7). (J) Relative mitochondrial membrane potential was measured in shSLC25A51 or shGFP expressing AML cells by flow cytometry using MitoTracker™ Red CMXRos staining. Data shown as mean ± SD (n=3-6). *p* values in this figure were determined by unpaired, two-tailed Student’s t-test (for two groups) or ANOVA with postdoc Dunnett’s multiple comparisons analysis (for groups of three). \**p* <0.05, \*\**p* <0.01, \*\*\**p* <0.001, and \*\*\*\**p* <0.0001 versus control.

**Table S2.** Proliferation rate upon depletion of SLC25A51 in human myeloid cells.

| Cell line | Proliferation rate (Mean ± SEM) | |
| --- | --- | --- |
|  | shGFP | shSLC25A51 |
| MOLM13 | 1.215 ± 0.012 | 0.013 ± 0.007**** |
| MOLM14 | 1.003 ± 0.014 | 0.220 ± 0.007**** |
| U937 | 1.261 ± 0.069 | 0.841 ± 0.089** |
| MV411 | 1.021 ± 0.069 | 0.757 ± 0.083* |
| THP-1 | 0.898 ± 0.042 | 0.496 ± 0.132* |
| NB4 | 1.223 ± 0.003 | 0.655 ± 0.006**** |
| Kasumi-1 | 0.648 ± 0.006 | 0.374 ± 0.005*** |
| HAL01 | 0.599 ± 0.021 | -0.012 ± 0.006*** |
| MDSL | 0.815 ± 0.002 | 0.663 ± 0.014** |
| TF-1 | 1.211 ± 0.006 | 0.690 ± 0.009**** |

*p* values were determined by the unpaired, two-tailed Student's *t*-test. Data shown as mean ± SEM. \**p* < 0.05, \*\**p* < 0.01, \*\*\**p* < 0.001, and \*\*\*\**p* < 0.0001 versus control
Proliferation rate was determined using the following formula:
Proliferation Rate (Doublings per day) = $\log_2$ (Final cell count (D5)/Initial cell count (D0))/5 (days)

**Figure S3.**
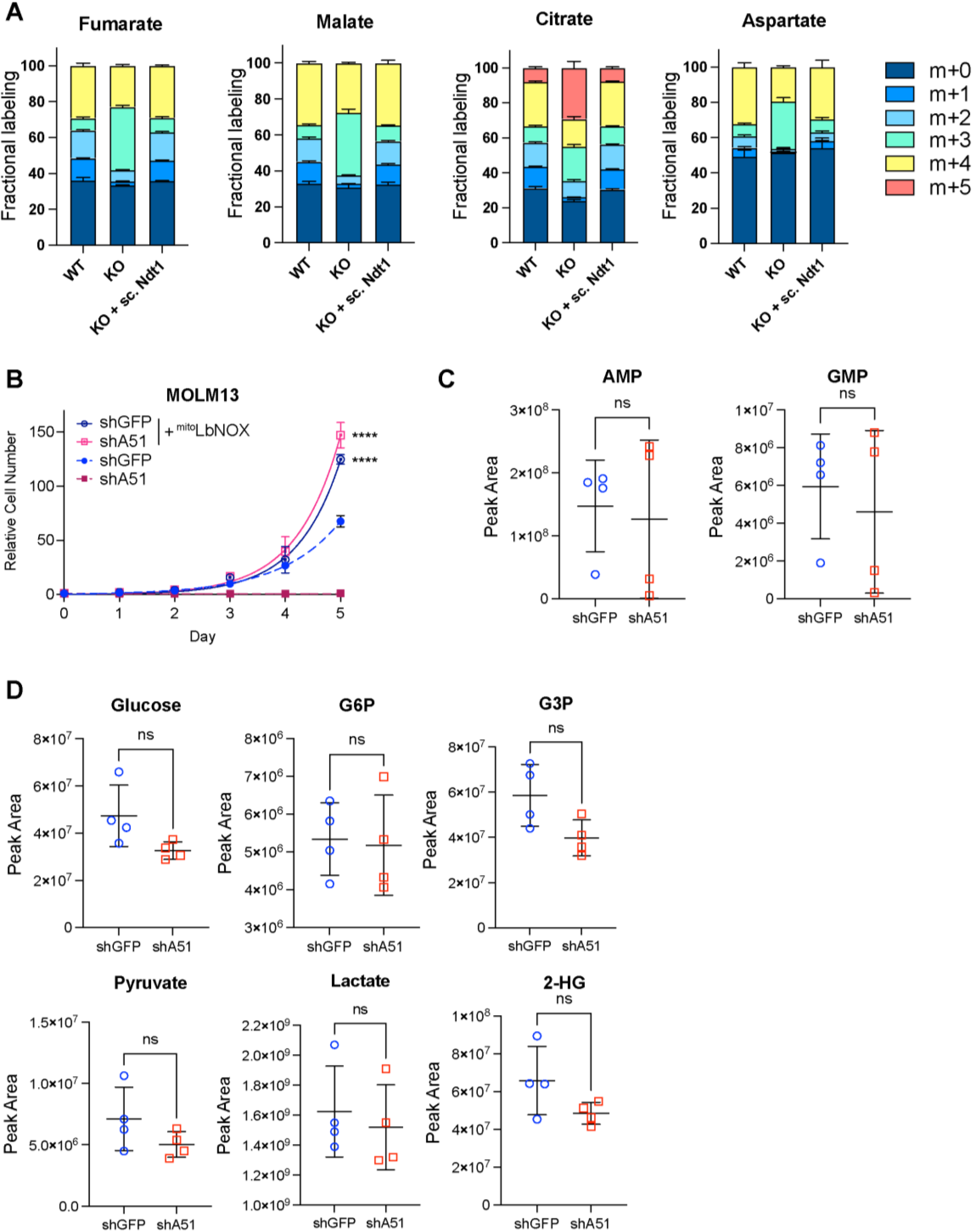
Loss of SLC25A51 rewires glutamine metabolism. (A) Fractional labeling of fumarate, malate, aspartate, and citrate derived from [^13^C_5_,^15^N_2_]-glutamine; mean ± SD (n=4). (B) Growth curves of shSLC25A51 or shGFP expressing MOLM-13 cells in the presence or absence of ^mito^*Lb*NOX. Data shown as mean ± SD (n=3). *p* values were determined by unpaired, two-tailed Student’s t-test at end point. \*\*\*\**p* <0.0001 versus the absence of ^mito^*Lb*NOX. (C) Total cellular levels of AMP and GMP in shSLC25A51 and shGFP expressing U937 cells, measured by UHPLC-MS/MS. Data shown as mean ± SD (n=4). (D) Total cellular levels of glucose, glucose-6-phosphate (G6P), glyceraldehyde-3-phosphate (G3P), pyruvate, lactate, and 2-hydroxyglutarate (2-HG) in shSLC25A51 and shGFP expressing U937 cells were measured by UHPLC-MS/MS. Data shown as mean ± SD (n=4). *p* values in C and D were determined by unpaired, two-tailed Student’s t-test.

**Figure S4.**
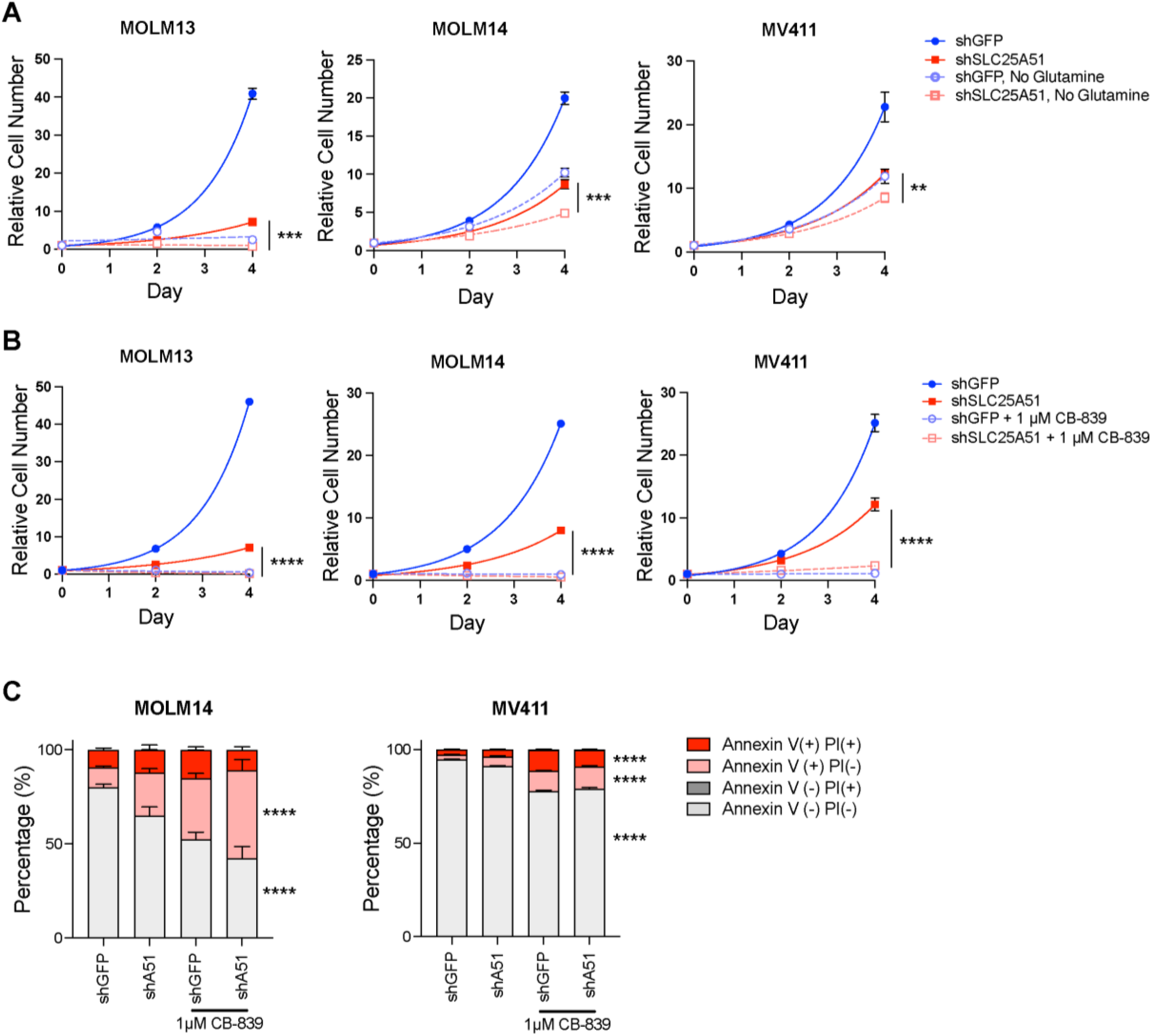
Depletion of SLC25A51 sensitizes AML cells to glutamine deprivation and glutaminase inhibitor. (A) Growth curves of shSLC25A51 or shGFP expressing AML cells grown with indicated concentrations of glutamine in the media. Cell counts were measured at indicated time using flow cytometry. Data shown as mean ± SD (n=3). (B) Growth curves of shSLC25A51 or shGFP expressing AML cells treated with 1 µM CB-839. Cell counts were measured at indicated times using flow cytometry. Data shown as mean ± SD (n=3). (C) Apoptotic rates of shSLC25A51 or shGFP expressing AML cells treated with 1 µM CB-839 and measured by annexin V-FITC and Propidium Iodide (PI) staining. Bar graph quantifying the percentage of living, early-stage apoptotic, late-stage apoptotic, and dead cells. Data shown as mean ± SD (n=3). *p* values were determined by unpaired, two-tailed Student’s t-test at end point. \*\**p* <0.01, \*\*\**p* <0.001, and \*\*\*\**p* <0.0001 versus shSLC25A51 group.

**Table S3.**
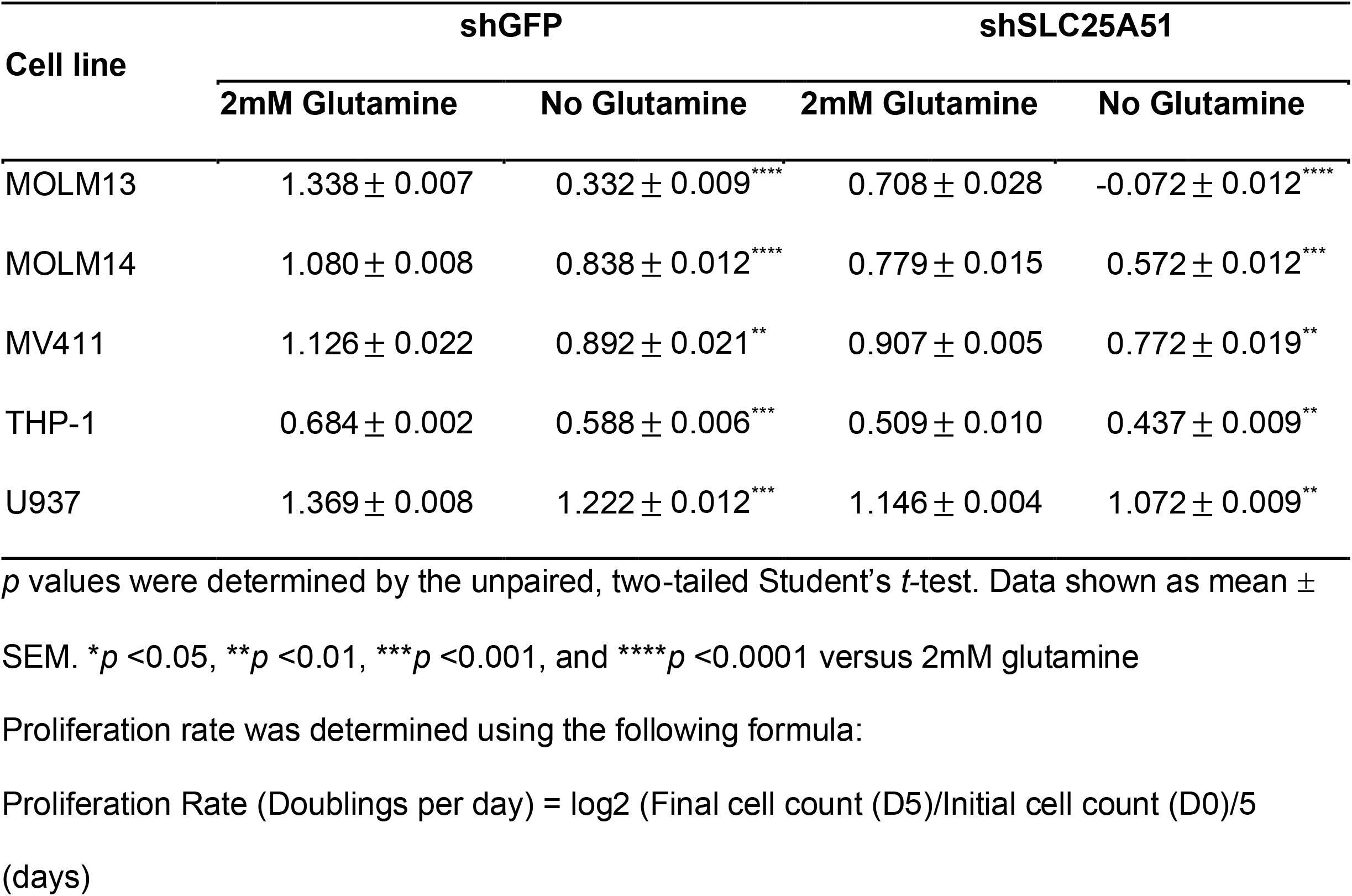
Proliferation rate of SLC25A51 depleted cell upon glutamine removal.

**Table S4.** Proliferation rate of SLC25A51 depleted cell upon CB-839 treatment.

| Cell line | shGFP |  | shSLC25A51 |  |
| --- | --- | --- | --- | --- |
|  | control | 1uM CB-839 | control | 1uM CB-839 |
| MOLM13 | 1.381 ± 0.001 | -0.372 ± 0.030**** | 0.707 ± 0.006 | -0.702 ± 0.014**** |
| MOLM14 | 1.162 ± 0.004 | -0.036 ± 0.035**** | 0.749 ± 0.001 | -0.214 ± 0.022**** |
| MV411 | 1.163 ± 0.012 | 0.031 ± 0.050**** | 0.899 ± 0.018 | 0.301 ± 0.021**** |
| THP-1 | 0.944 ± 0.009 | 0.806 ± 0.003*** | 0.683 ± 0.004 | 0.575 ± 0.008*** |
| U937 | 1.404 ± 0.008 | 1.037 ± 0.019**** | 0.917 ± 0.008 | 0.702 ± 0.004**** |

*p* values were determined by the unpaired, two-tailed Student's *t*-test. Data shown as mean ±
SEM. \**p* <0.05, \*\**p* <0.01, \*\*\**p* <0.001, and \*\*\*\**p* <0.0001 versus control
Proliferation rate was determined using the following formula:
Proliferation Rate (Doublings per day) = $\log_2$ (Final cell count (D5)/Initial cell count (D0))/5 (days)

## References

1. Panina SB, Pei J, Baran N, Konopleva M, Kirienko NV. Utilizing Synergistic Potential of Mitochondria-Targeting Drugs for Leukemia Therapy. Front Oncol. 2020;10:435.

2. Molina JR, Sun Y, Protopopova M, Gera S, Bandi M, Bristow C, et al. An inhibitor of oxidative phosphorylation exploits cancer vulnerability. Nat Med. 2018;24:1036–46.

3. Matre P, Velez J, Jacamo R, Qi Y, Su X, Cai T, et al. Inhibiting glutaminase in acute myeloid leukemia: metabolic dependency of selected AML subtypes. Oncotarget. 2016;7:79722–35.

4. Pachnis P, Wu Z, Faubert B, Tasdogan A, Gu W, Shelton S, et al. In vivo isotope tracing reveals a requirement for the electron transport chain in glucose and glutamine metabolism by tumors. Sci Adv. 2022;8:eabn9550.

5. Thein MS, Ershler WB, Jemal A, Yates JW, Baer MR. Outcome of older patients with acute myeloid leukemia: An Analysis of SEER Data Over 3 Decades. Cancer. 2013;119:2720–7.

6. Cambronne XA, Kraus WL. Location, Location, Location: Compartmentalization of NAD+ Synthesis and Functions in Mammalian Cells. Trends in Biochemical Sciences. 2020;45:858–73.

7. Cambronne XA, Stewart ML, Kim D, Jones-Brunette AM, Morgan RK, Farrens DL, et al. Biosensor reveals multiple sources for mitochondrial NAD ^+^. Science. 2016;352:1474–7.

8. Vazquez BN, Fernández-Duran I, Vaquero A. Sirtuins in hematopoiesis and blood malignancies. Sirtuin Biology in Medicine. Elsevier; 2021. page 373–91.

9. Farber M, Chen Y, Arnold L, Möllmann M, Boog-Whiteside E, Lin Y-A, et al. Targeting CD38 in acute myeloid leukemia interferes with leukemia trafficking and induces phagocytosis. Sci Rep. 2021;11:22062.

10. Luongo TS, Eller JM, Lu M-J, Niere M, Raith F, Perry C, et al. SLC25A51 is a mammalian mitochondrial NAD+ transporter. Nature. 2020;588:174–9.

11. Kory N, uit de Bos J, van der Rijt S, Jankovic N, Güra M, Arp N, et al. MCART1/SLC25A51 is required for mitochondrial NAD transport. SciAdv. 2020;eabe5310.

12. Girardi E, Agrimi G, Goldmann U, Fiume G, Lindinger S, Sedlyarov V, et al. Epistasis-driven identification of SLC25A51 as a regulator of human mitochondrial NAD import. Nat Commun. 2020;11:6145.

13. Jung N, Dai B, Gentles AJ, Majeti R, Feinberg AP. An LSC epigenetic signature is largely mutation independent and implicates the HOXA cluster in AML pathogenesis. Nat Commun. 2015;6:8489.

14. Metzeler KH, Hummel M, Bloomfield CD, Spiekermann K, Braess J, Sauerland M-C, et al. An 86-probe-set gene-expression signature predicts survival in cytogenetically normal acute myeloid leukemia. Blood. 2008;112:4193–201.

15. Li Z, Herold T, He C, Valk PJM, Chen P, Jurinovic V, et al. Identification of a 24-Gene Prognostic Signature That Improves the European LeukemiaNet Risk Classification of Acute Myeloid Leukemia: An International Collaborative Study. JCO. 2013;31:1172–81.

16. Panina SB, Pei J, Kirienko NV. Mitochondrial metabolism as a target for acute myeloid leukemia treatment. Cancer Metab. 2021;9:17.

17. de Beauchamp L, Himonas E, Helgason GV. Mitochondrial metabolism as a potential therapeutic target in myeloid leukaemia. Leukemia. 2022;36:1–12.

18. Todisco S, Agrimi G, Castegna A, Palmieri F. Identification of the Mitochondrial NAD+ Transporter in Saccharomyces cerevisiae. Journal of Biological Chemistry. 2006;281:1524–31.

19. Pandey R, Riley CL, Mills EM, Tiziani S. Highly sensitive and selective determination of redox states of coenzymes Q9 and Q10 in mice tissues: Application of orbitrap mass spectrometry. Analytica Chimica Acta. 2018;1011:68–76.

20. Gallipoli P, Giotopoulos G, Tzelepis K, Costa ASH, Vohra S, Medina-Perez P, et al. Glutaminolysis is a metabolic dependency in FLT3ITD acute myeloid leukemia unmasked by FLT3 tyrosine kinase inhibition. Blood. 2018;131:1639–53.

21. Gregory MA, Nemkov T, Park HJ, Zaberezhnyy V, Gehrke S, Adane B, et al. Targeting Glutamine Metabolism and Redox State for Leukemia Therapy. Clin Cancer Res. 2019;25:4079–90.

22. Alkan HF, Bogner-Strauss JG. Maintaining cytosolic aspartate levels is a major function of the TCA cycle in proliferating cells. Molecular & Cellular Oncology. 2019;6:e1536843.

23. Birsoy K, Wang T, Chen WW, Freinkman E, Abu-Remaileh M, Sabatini DM. An Essential Role of the Mitochondrial Electron Transport Chain in Cell Proliferation Is to Enable Aspartate Synthesis. Cell. 2015;162:540–51.

24. Sullivan LB, Gui DY, Hosios AM, Bush LN, Freinkman E, Vander Heiden MG. Supporting Aspartate Biosynthesis Is an Essential Function of Respiration in Proliferating Cells. Cell. 2015;162:552–63.

25. Titov DV, Cracan V, Goodman RP, Peng J, Grabarek Z, Mootha VK. Complementation of mitochondrial electron transport chain by manipulation of the NAD+/NADH ratio. Science. 2016;352:231–5.

26. Sallin O, Reymond L, Gondrand C, Raith F, Koch B, Johnsson K. Semisynthetic biosensors for mapping cellular concentrations of nicotinamide adenine dinucleotides. eLife. 2018;7:e32638.

27. Fu Z, Kim H, Morse PT, Lu M-J, Hüttemann M, Cambronne XA, et al. The mitochondrial NAD+ transporter SLC25A51 is a fasting-induced gene affecting SIRT3 functions. Metabolism. 2022;135:155275.

28. Stevens BM, Jones CL, Pollyea DA, Culp-Hill R, D’Alessandro A, Winters A, et al. Fatty acid metabolism underlies venetoclax resistance in acute myeloid leukemia stem cells. Nat Cancer. 2020;1:1176–87.

29. Yan D, Franzini A, Pomicter AD, Halverson BJ, Antelope O, Mason CC, et al. SIRT5 Is a Druggable Metabolic Vulnerability in Acute Myeloid Leukemia. Blood Cancer Discov. 2021;2:266–87.

30. Chowdhry S, Zanca C, Rajkumar U, Koga T, Diao Y, Raviram R, et al. NAD metabolic dependency in cancer is shaped by gene amplification and enhancer remodelling. Nature. 2019;569:570–5.

31. Chedere A, Mishra M, Kulkarni O, Sriraman S, Chandra N. Personalized quantitative models of NAD metabolism in hepatocellular carcinoma identify a subgroup with poor prognosis. Front Oncol. 2022;12:954512.

32. Maric T, Bazhin A, Khodakivskyi P, Mikhaylov G, Solodnikova E, Yevtodiyenko A, et al. A bioluminescent-based probe for in vivo non-invasive monitoring of nicotinamide riboside uptake reveals a link between metastasis and NAD+ metabolism. Biosensors and Bioelectronics. 2023;220:114826.

33. Lagadinou ED, Sach A, Callahan K, Rossi RM, Neering SJ, Minhajuddin M, et al. BCL-2 Inhibition Targets Oxidative Phosphorylation and Selectively Eradicates Quiescent Human Leukemia Stem Cells. Cell Stem Cell. 2013;12:329–41.

34. Baccelli I, Gareau Y, Lehnertz B, Gingras S, Spinella J-F, Corneau S, et al. Mubritinib Targets the Electron Transport Chain Complex I and Reveals the Landscape of OXPHOS Dependency in Acute Myeloid Leukemia. Cancer Cell. 2019;36:84-99.e8.

35. Subedi A, Liu Q, Ayyathan DM, Sharon D, Cathelin S, Hosseini M, et al. Nicotinamide phosphoribosyltransferase inhibitors selectively induce apoptosis of AML stem cells by disrupting lipid homeostasis. Cell Stem Cell. 2021;28:1851-1867.e8.

36. Jones CL, Stevens BM, Pollyea DA, Culp-Hill R, Reisz JA, Nemkov T, et al. Nicotinamide Metabolism Mediates Resistance to Venetoclax in Relapsed Acute Myeloid Leukemia Stem Cells. Cell Stem Cell. 2020;

37. Bhingarkar A, Vangapandu HV, Rathod S, Hoshitsuki K, Fernandez CA. Amino Acid Metabolic Vulnerabilities in Acute and Chronic Myeloid Leukemias. Front Oncol. 2021;11:694526.

38. Lv H, Lv G, Chen C, Zong Q, Jiang G, Ye D, et al. NAD+ Metabolism Maintains Inducible PD-L1 Expression to Drive Tumor Immune Evasion. Cell Metabolism. 2021;33:110-127.e5.

39. Wang Y, Wang F, Wang L, Qiu S, Yao Y, Yan C, et al. NAD+ supplement potentiates tumor-killing function by rescuing defective TUB-mediated NAMPT transcription in tumor-infiltrated T cells. Cell Reports. 2021;36:109516.

40. Hamity MV, White SR, Walder RY, Schmidt MS, Brenner C, Hammond DL. Nicotinamide riboside, a form of vitamin B3 and NAD+ precursor, relieves the nociceptive and aversive dimensions of paclitaxel-induced peripheral neuropathy in female rats. Pain. 2017;158:962–72.

41. Liu H, Smith CB, Schmidt MS, Cambronne XA, Cohen MS, Migaud ME, et al. Pharmacological bypass of NAD ^+^salvage pathway protects neurons from chemotherapy-induced degeneration. Proc Natl Acad Sci USA. 2018;115:10654–9.

42. Achreja A, Yu T, Mittal A, Choppara S, Animasahun O, Nenwani M, et al. Metabolic collateral lethal target identification reveals MTHFD2 paralogue dependency in ovarian cancer. Nat Metab. 2022;4:1119–37.

43. Bai L, Yang Z-X, Ma P-F, Liu J-S, Wang D-S, Yu H-C. Overexpression of SLC25A51 promotes hepatocellular carcinoma progression by driving aerobic glycolysis through activation of SIRT5. Free Radical Biology and Medicine. 2022;182:11–22.

44. Wei P, Bott AJ, Cluntun AA, Morgan JT, Cunningham CN, Schell JC, et al. Mitochondrial pyruvate supports lymphoma proliferation by fueling a glutamate pyruvate transaminase 2–dependent glutaminolysis pathway. Sci Adv. 2022;8:eabq0117.

45. Baran N, Lodi A, Dhungana Y, Herbrich S, Collins M, Sweeney S, et al. Inhibition of mitochondrial complex I reverses NOTCH1-driven metabolic reprogramming in T-cell acute lymphoblastic leukemia. Nat Commun. 2022;13:2801.

46. Okuda S, Yamada T, Hamajima M, Itoh M, Katayama T, Bork P, et al. KEGG Atlas mapping for global analysis of metabolic pathways. Nucleic Acids Research. 2008;36:W423–6.

47. Wishart DS, Guo A, Oler E, Wang F, Anjum A, Peters H, et al. HMDB 5.0: the Human Metabolome Database for 2022. Nucleic Acids Research. 2022;50:D622–31.

